# Characterizing constitutive promoters across the Proteobacteria

**DOI:** 10.1101/2023.11.02.564426

**Authors:** Layla Schuster, Catalina Mejia, Lidimarie Trujillo Rodriguez, Emily Kairalla, Christopher R. Reisch, Marc G. Chevrette, Raquel Dias

## Abstract

Although research on promoters has spanned decades, the precise prediction of promoter activity from DNA sequence remains a challenge even in model organisms. Recent literature has identified important differences in the core sequence of σ^70^ promoters across classes of Proteobacteria as well as a lack of transferability when promoters are moved from host to host. Currently, there is a need for synthetic constitutive promoters spanning a range of expression levels in species outside of *Escherichia coli.* Additionally, characterization data defining behavior of the same promoter across multiple species would be extremely valuable to the field. Here, we analyzed promoter activity in three classes of Proteobacteria, which enabled us to better understand the sequence elements correlated with a strong promoter in different hosts. In doing so, we identified and characterized constitutive promoters spanning a range of expression in these species for community use and described the portability of a subset of these promoters as they were moved between hosts. These promoter libraries have broad applications as predictable genetic tools to control gene expression in diverse species (1–3). This work adds to the toolkit for gene expression in non-model bacteria and is a step towards the larger goal of accurate promoter prediction in a given host from a *de novo* sequence.

## Introduction

Promoters are essential regulators of gene expression. They are largely responsible for the modulation of cellular responses to various stimuli and affect organism behavior by mediating rates of transcriptional initiation (4–6). As such, they have immense utility as a genetic tool to control gene expression (7–9). Bacterial transcription requires the tight association of the RNA polymerase to a σ specificity factor. In Gram-negative bacteria, σ^70^ is responsible for the transcription of most active promoters during log-phase growth (10–14). The σ^70^ proteins are modular, with three highly conserved domains (σ domain 2 (σD2), σD3 and σD4) and an amino-terminal domain that is poorly conserved (σD1) (15, 16). Promoters regulated by σ^70^ are generally divided into discrete sequence elements including the –10 and –35 hexamers, comprising the core promoter, and the UP element, spacer region, and extended –10 element, if present (16–20). These elements interact specifically with domains of σ^70^ during promoter binding (15, 21). Region 2.4 of σD2 directly interacts with the –10 element and has nearly 100% identity in Proteobacteria while region 4.2 of σD4 interacts with the –35 element and is less but still very well conserved (20–26). Despite decades of research on promoters and transcriptional initiation (10, 11), we are still unable to predict the activity of a promoter from its sequence and how that activity will vary when the promoter is moved between hosts (5, 27). More urgently, constitutive promoters that are characterized outside of *E. coli* and validated in different species are severely lacking in the field.

The closer core promoter sequences are to the consensus, the stronger the promoter (28). In *E. coli*, a σ^70^-dependent consensus promoter sequence includes a 5′ – TTGACA – 3′ –35 hexamer and 5′ – TATAAT – 3′ –10 hexamer separated by a 17 bp spacer (29). For non-model species in some classes of Proteobacteria, the consensus promoter sequence has been predicted to be very similar to that of *E. coli*, but studies have conflicting results (30, 31). Accordingly, studies have also shown inconsistencies in expression levels when the same promoter is moved between Proteobacteria. For example, *E. coli* promoters retain similar relative activity levels in some *Pseudomonas* and Alphaproteobacteria species but the reverse is not true in either case (30, 32–36). This is known as the transcriptional laxity phenomenon (37), with some species able to recognize promoters with more laxity than others. Without characterized promoters in these Proteobacteria, researchers are constrained to genetic tools optimized for *E. coli* or moving systems into *E. coli* even when it’s not the most ideal host.

In the work described here, we reutilize a previously described *E. coli* constitutive promoter toolbox to test promoters in 15 species across the Alpha-, Beta-, and Gammaproteobacteria. We present characterized libraries of 15 to 43 promoter sequences for each species with expression spanning 3-5 orders of magnitude within each library. We then surveyed the promoter libraries within and across species to identify conserved elements and other characteristics of high expressing promoters. Specifically, we compared sequences of the core hexamers, the presence or absence of an extended –10 element, and the GC content of the spacer region. Finally, we tested the transferability of a subset of promoters from the libraries by rescreening them in four species across the three Proteobacterial classes.

## Materials and Methods

### Library Construction and Transformations

This work utilized a previously described synthetic constitutive promoter library with 4350 unique promoter variants, generously provided by the Salis lab (38). To move the promoter variants into our previously developed broad-host-range vectors, we followed our protocol for new part addition in the combinatorial assembly workflow (2). Namely, the promoter variant and *mRFP* reporter gene were amplified as a single piece from the promoter library via PCR, treated with Dpn1 restriction enzyme, run on a gel for size verification, and gel purified. The same protocol was followed for amplification of the plasmid backbone and amplified parts were assembled using NEB HiFi Assembly. Here, the backbone contained a pBBR origin and gentamicin marker for all species except for *Acinetobacter baylyi*, where the RK2 origin was used. Recipient strains were transformed via electroporation, conjugation, or natural transformation, as specified in Table 3.

#### Electroporation

Cells to be made electrocompetent were taken either from overnight cultures or subcultured from an overnight growth and made electrocompetent when in mid-log phase. The protocol for the preparation of electrocompetent cells was as follows: 6 mL of each wild-type strain was incubated with shaking in fresh media with the specific culturing conditions listed in Table 2. The total culture was then spun at 5000 rpm for 2 min to pellet the cells. Culture supernatants were aspirated, cell pellets resuspended in 1 mL 300 mM sucrose at room temperature, and then centrifuged again for 2 min at 5000 rpm. This process was repeated to wash the cell pellet with sucrose twice and then the pellet was resuspended in a final volume of 1:10 of the initial culture volume. 50 µL of each sucrose-cell suspension was then transferred to a 1-mm-gap-width electroporation cuvette and cells were electroporated at the specified voltage for each strain (Table 3). Cells were recovered in 1 mL of their respective recovery media and incubated for 2 h in a deep-well plate with the specified conditions (Table 2) before plating on to selection plates.

**Table 1.** Strains investigated in this study.

| Strain | Phylogenetic class |
| --- | --- |
| *Acinetobacter baylyi* ADP1 | Gammaproteobacteria |
| *Agrobacterium fabrum* C58 | Alphaproteobacteria |
| *Burkholderia thailandensis* E264* | Betaproteobacteria |
| DSM *Massilia niabensis* | Betaproteobacteria |
| *Ensifer meliloti* SM1021 | Alphaproteobacteria |
| *Enterobacter cloacae* ATCC 13047 | Gammaproteobacteria |
| *Escherichia coli* MG1655 | Gammaproteobacteria |
| *Pseudomonas aeruginosa* PAO1 | Gammaproteobacteria |
| *Pseudomonas putida* KT2440 | Gammaproteobacteria |
| *Pseudomonas syringae* pv. *syringae* IBSBF 281 | Gammaproteobacteria |
| *Rhodopseudomonas palustris* | Alphaproteobacteria |
| *Ruegeria* sp. TM1040 | Alphaproteobacteria |
| *Salmonella enterica* Typhimurium | Gammaproteobacteria |
| *Sulfitobacter* sp. EE-36* | Alphaproteobacteria |
| *Xanthomonas campestris* ATCC 33913 | Gammaproteobacteria |

\* Strains included in promoter swapping experiments

**Table 2.** Strain growth conditions.

| Strain | Growth medium | Incubation temperature | Gentamicin concentration |
| --- | --- | --- | --- |
| *Acinetobacter baylyi* ADP1 | LB | 30°C | 20 µg/mL |
| *Agrobacterium fabrum* C58 | LB | 30°C | 100 µg/mL |
| *Burkholderia thailandensis* E264 | LSLB | 37°C | 100 µg/mL |
| DSM *Massilia niabensis* | R2A | 30°C | 5 µg/mL |
| *Ensifer meliloti* SM1021 | LB | 30°C | 100 µg/mL |
| *Enterobacter cloacae* ATCC 13047 | LB | 37°C | 20 µg/mL |
| *Escherichia coli* MG1655 | LB | 37°C | 20 µg/mL |
| *Pseudomonas aeruginosa* PAO1 | LB | 37°C | 100 µg/mL |
| *Pseudomonas putida* KT2440 | LB | 30°C | 100 µg/mL |
| *Pseudomonas syringae* pv. *syringae* IBSBF 281 | LB | RT | 100 µg/mL |
| *Rhodopseudomonas palustris* | LB | 30°C | 100 µg/mL |
| *Ruegeria* sp. TM1040 | ½ YTSS | 30°C | 50 µg/mL |
| *Salmonella enterica* Typhimurium | LB | 37°C | 100 µg/mL |
| *Sulfitobacter* sp. EE-36 | ½ YTSS | 30°C | 20 µg/mL |
| *Vibrio vulnificus* | LB | 30°C | 100 µg/mL |
| *Xanthomonas campestris* ATCC 33913 | LB | 30°C | 20 µg/mL |

**Table 3.** Transformation conditions.

| Strain | Transformation method | Recovery medium |
| --- | --- | --- |
| *Acinetobacter baylyi* ADP1 | Natural transformation | LB |
| *Agrobacterium fabrum* C58 | Electroporation<sup>2‡</sup> | LB |
| *Burkholderia thailandensis* E264 | Electroporation<sup>1‡</sup> | LSLB |
| DSM *Massilia niabensis* | Conjugation | R2A |
| *Ensifer meliloti* SM1021 | Conjugation | LB |
| *Enterobacter cloacae* ATCC 13047 | Electroporation<sup>2†</sup> | LB |
| *Escherichia coli* MG1655 | Electroporation<sup>2†</sup> | LB |
| *Pseudomonas aeruginosa* PAO1 | Electroporation<sup>1†</sup> | LB |
| *Pseudomonas putida* KT2440 | Electroporation<sup>1†</sup> | LB |
| *Pseudomonas syringae* pv. *syringae* IBSBF 281 | Electroporation<sup>1†</sup> | LB |
| *Rhodopseudomonas palustris* | Conjugation | LB |
| *Ruegeria* sp. TM1040 | Electroporation<sup>2‡</sup> | ½ YTSS |
| *Salmonella enterica* Typhimurium | Electroporation<sup>2†</sup> | LB |
| *Sulfitobacter* sp. EE-36 | Electroporation<sup>2‡</sup> | ½ YTSS |
| *Xanthomonas campestris* ATCC 33913 | Electroporation<sup>2†</sup> | LB |

<sup>1</sup> Cells made electrocompetent taken from an overnight. growth
<sup>2</sup> Cells were subcultured from an overnight growth and made electrocompetent when in mid-log phase
† Electroporated with 1.8 kV, 1 pulse
‡ Electroporated with 2.2 kV, 1 pulse

#### Natural transformation

A natural transformation protocol was followed for *A. baylyi* which was adapted from a previous protocol (39). Here, 5 mL of fresh LB was inoculated with wild-type *A. baylyi* from a glycerol stock and grown overnight at 30°C. The next day, 1 mL of fresh LB was inoculated with 70 µL of this culture and approximately 100 ng of the plasmid were incubated for 3 h before plating onto selection plates.

#### Conjugation

Conjugation was performed using the RP4 system. On the day prior to conjugation, 5 mL cultures were inoculated from glycerol stocks of wild-type strains, all requiring donor strains of *E. coli,* and an *E. coli* helper strain containing pEVS104, and grown overnight. The following day, donor and helper cultures were spun down separately at 10 000 rpm for 1 min and resuspended in fresh media to remove residual antibiotic. A sufficient volume of donor and helper cultures was pelleted and resuspended such that each conjugation used 500 µL of both donor and helper strains in addition to 500 µL of the recipient strain to be transformed. Each mixture of donor, helper, and recipient was centrifuged at 10 000 rpm for 1 min and the supernatant decanted, leaving approximately 100 µL of media to resuspend the pellet. The resuspensions were spotted on agar plates and incubated on the benchtop overnight. Each spot was streaked on to selection plates the following day.

### Library Picking

For the promoter library in each strain, colonies were picked to inoculate 92 wells of a 96-well ‘picked’ plate, with 2 wells left as media blanks and 2 wells inoculated with the wild-type strain to be used as background fluorescence controls. The transformants were picked by hand and a mix of red, pink, and white colonies were chosen to provide a range of promoter expression levels in the sampling. The picked plate was incubated overnight with shaking and the following day, used to inoculate four 96-well plates, including screening plates in triplicate and a stock plate. Stock and screening plates were inoculated at 1:200 for fast-growing strains or 1:50 for slow-growing strains. The stock plate was incubated until cultures were in stationary phase, glycerol was added to a final concentration of 25%, and then the plate was frozen at –80°C to be used for library strain retrieval. The screening plates (Costar, black, clear-bottom) were used to measure mRFP expression. After inoculation of the stock and screening plates, the picked plate was frozen to be used for downstream PCR amplification.

### Library Screening

Fluorescence readings of mRFP were taken to measure promoter variant activity. Inoculation of the screening plates from the picked plate marked the start of the screen. OD_660_ and fluorescence readings were taken in a plate reader (Molecular Devices SpectraMax M3) every hour from 0-5 hours for faster growing strains, or from 4-8 hours for slower growing strains, and a final timepoint was taken at 24 hours. Absorbance at 660 nm is used to measure growth as 600 nm is significantly absorbed by mRFP (40). Libraries were screened in triplicate.

### Promoter Mapping

To determine the promoter sequence associated with the fluorescence measurement of each well of the screening plates, a hierarchical barcoding scheme was employed. Here, unique pairings of barcoded primers were assigned to each well of the screening plate to amplify the promoter region and in a second PCR reaction, unique indexing primers were used to differentiate screening plates, including replicate plates of the same species. Specifically, primers were aliquoted into wells of 96-well PCR plates according to predetermined maps such that each well has a unique combination of forward and reverse primers. These aliquoted primer plates were prepared in bulk and frozen ahead of experiments for increased efficiency. To prepare the PCR reaction, OneTaq master mix was added to the primer-aliquoted PCR plates and cell material was added to the wells from the thawed picked plate via plate stamper. The promoter variant part was amplified with 30 cycles, the reaction products from each plate were pooled into a single tube, and a small volume was run on a gel for verification. 200 µl of the remaining product was PCR purified and 6 μL was used in the second PCR where indexing primers and adapters were added for Illumina sequencing. Here, amplification was limited to 7-10 cycles. Pooled plate amplification products were gel purified, measured with Qubit, and sequenced using 2 x 150 reads on a shared Novaseq6000 instrument. In this way, each promoter variant in every test species could be identified and mapped to a fluorescence measurement characterizing its behavior. Plates were sequenced in technical duplicates except for *A. fabrum*, *E. coli*, *P. putida*, and *R*. sp. TM1040, which did not have sequencing replicates.

### Data Analysis

All calculations and data analysis were performed using Microsoft Excel and custom R scripts to parse promoter sequences into core promoter, spacer, UP, and extended −10 elements. For library screens in each bacterial strain, absorbance and fluorescence data were organized by timepoint. Optical density was adjusted to a 1 cm pathlength by dividing by a factor of 0.56 for a culture volume of 200 µL in the wells of the 96-well plate, as done in a previous study (2). The average raw fluorescence data of the replicates was used directly in the graphs and analysis included herein unless specified otherwise.

Sequencing data for each species was analyzed through a pipeline that included joining the reads with Flash, trimming the reads with Cutadapt, and performing quality checks with Fastqc. Expression from promoters of the entire screened library in each species is represented in Figure 1. A subset of the entire promoter library passed quality checking and those sequences are analyzed in depth in this work. Only reads that were consistent across the two replicate sequencing runs were included in the final promoter set. For the four species without replicate sequencing data (*E. coli*, *A. fabrum*, *P. putida*, and *R.* sp. TM1040), only wells with >30% of the total sequences having a single defined promoter sequence were included in the study. The original *in silico* design of the nonrepetitive promoters specifies a length of 78 bp (38). As such, promoters were separated into those with a length of 78 bp and those that deviated from that length. Promoters with a length of 78 bp are listed in Object 1 and were used to generate Weblogos and KpLogos as discussed below. Promoters with this length make up most of the library within each species. Promoters deviating from 78 bp are discussed separately as the core promoter elements are less positionally defined but nonetheless may still be highly active in some species.

**Figure 1.**
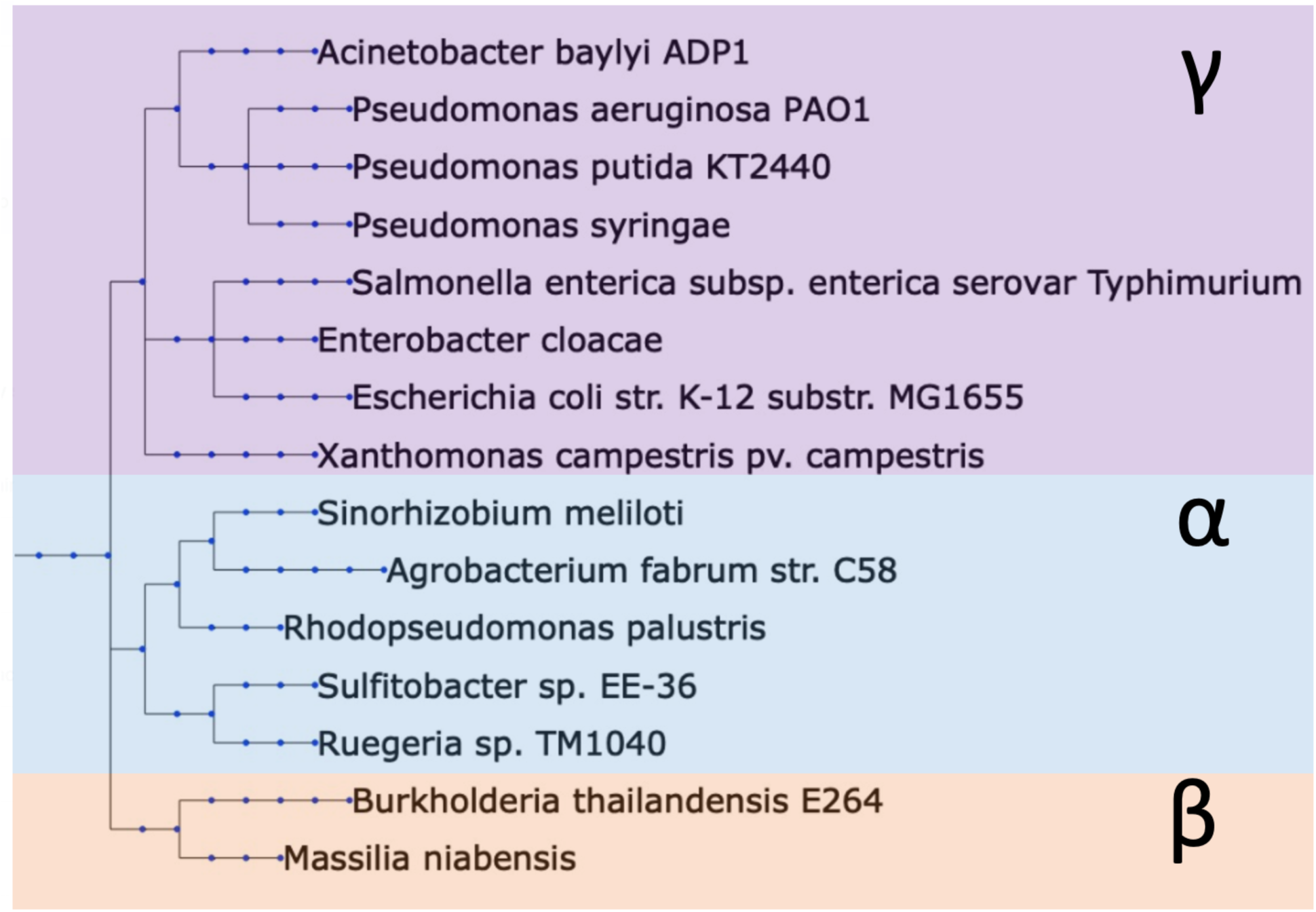
Phylogenetic tree of species included in the study. This study includes species from the Alpha-, Beta-, and Gammaproteobacteria classes. Ensifer meliloti referred to by its synonym Sinorhizobium meliloti.

Logos for core −10 and −35 elements and spacer elements were built using Weblogo at http://weblogo.berkeley.edu (41) and KpLogo probability logos (42). For all species except for *R.* sp. TM1040, promoters with a length of 78 bp were grouped into high, medium, and low activity promoters and logos were built for each group. In *R.* sp. TM1040, promoters were split into only high and low activity groups. Most promoter groups are separated by an order of magnitude of RFU expression. The full promoter sequences are included in a larger table in Object 1. Here, promoters are additionally separated into their discrete regions including the spacer, UP, extended −10, −10, and −35 elements.

### Promoter Transferability Experiments

Promoter variants were moved from source species, where library expression data was gathered, into target species, where promoter variants were measured again in a new host. Based on the mRFP expression data, promoter variants across a range of expression levels were chosen from four of the 15 species for transferability experiments, specifically *E. coli*, *P. aeruginosa*, *S.* sp. EE-36, and *B. thailandensis*. The promoter variants were struck from their associated wells in the stock plates and isolated colonies were grown in 5 mL cultures for plasmid extraction. The plasmids were then transformed into *E. coli* NEB5α and the transformants were grown, plasmids extracted, and the promoter region was sequenced via Sanger sequencing to verify that sequences matched Illumina results. Once verified, 11-12 plasmids from each source species were transformed into the target strains. The transformants were used to start 1 mL cultures in a deep well plate, incubated overnight, and subcultured 1:200 or 1:50 into screening plates for mRFP readings. Promoter transferability fluorescence measurements followed the same steps as library fluorescence measurements, with readings taken in exponential and late stationary phase.

## Results

### Constitutive Promoter Library Screens

A previously developed library of 4350 synthetic promoters, generously provided by the Salis lab, was utilized in this study (38). The library is a product of the Nonrepetitive Parts Calculator which generates nonrepetitive DNA sequences with the goal of increasing stability in constructed genetic systems. To characterize promoter activity in the Proteobacteria, we tested the libraries in each of the 15 species listed in Table 1, which are spread across the Alpha-, Beta-, and Gammaproteobacteria (Figure 1). We measured relative fluorescence units (RFU) of mRFP expression as a proxy for promoter activity and the promoters were sequenced by high-throughput sequencing. Approximately 100 library transformants were picked in each species and fluorescence and optical density readings were taken through exponential and late stationary phase of growth to measure promoter activity. The RFU output of all the promoters screened in each species are represented in Figure 2. The promoters were then amplified from each library transformant in a high-throughput workflow and sent for sequencing. To match promoter sequence to RFU output, the promoters were barcoded during PCR amplification such that each barcode corresponded to a well in the 96-well screening plate. In this way, we generated sets of promoters with activity spanning at least 3 orders of magnitude in species across the Alpha-, Beta-, and Gammaproteobacteria. After data processing and quality checks, the number of unique promoters included in each library ranged from 15 to 43 (Table 4). A comprehensive table of promoter sequences and outputs for all species included in this study are in Object 1.

**Figure 2.**
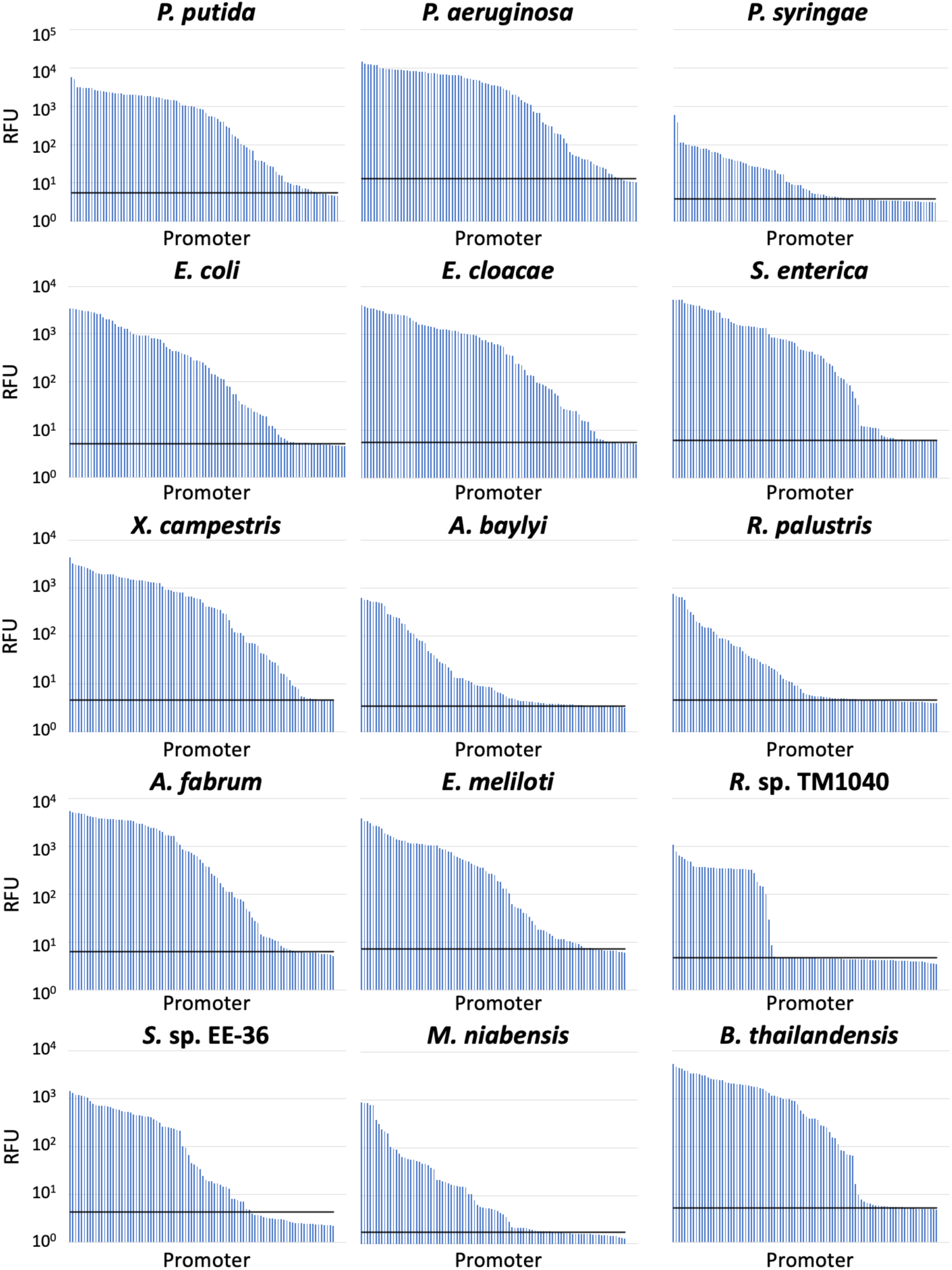
Range of expression from promoter libraries screened in each species. Data from late stationary phase of growth and includes 92 promoters in each library. Library picking described in Materials and Methods. Black horizontal line indicates fluorescence of wild type cells.

**Table 4.**
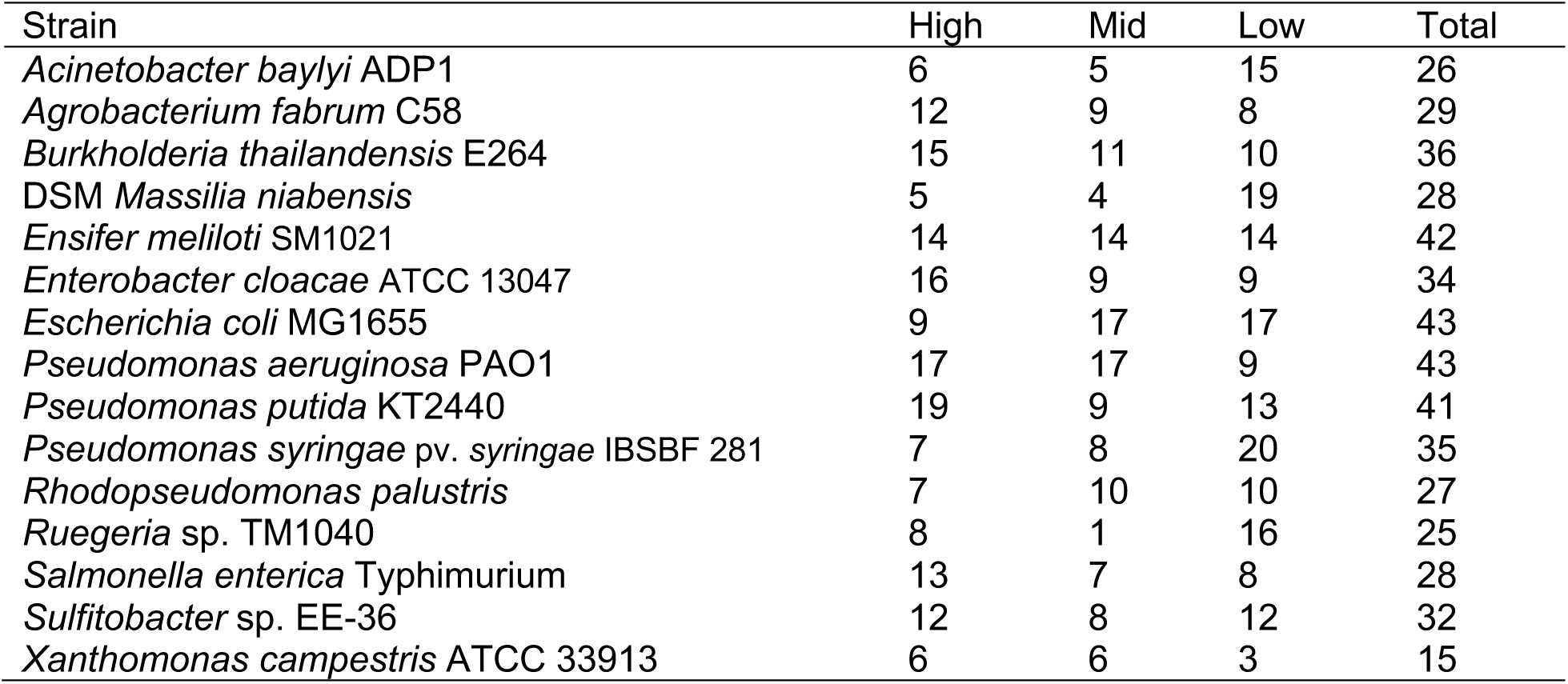
Promoter categorization. The number of promoters categorized as high, mid, and low expression sequences and total number of promoters in the library screened in each species is listed below. The full sequences of promoters in each category available in Object 1.

| Strain | High | Mid | Low | Total |
| --- | --- | --- | --- | --- |
| *Acinetobacter baylyi* ADP1 | 6 | 5 | 15 | 26 |
| *Agrobacterium fabrum* C58 | 12 | 9 | 8 | 29 |
| *Burkholderia thailandensis* E264 | 15 | 11 | 10 | 36 |
| DSM *Massilia niabensis* | 5 | 4 | 19 | 28 |
| *Ensifer meliloti* SM1021 | 14 | 14 | 14 | 42 |
| *Enterobacter cloacae* ATCC 13047 | 16 | 9 | 9 | 34 |
| *Escherichia coli* MG1655 | 9 | 17 | 17 | 43 |
| *Pseudomonas aeruginosa* PAO1 | 17 | 17 | 9 | 43 |
| *Pseudomonas putida* KT2440 | 19 | 9 | 13 | 41 |
| *Pseudomonas syringae* pv. *syringae* IBSBF 281 | 7 | 8 | 20 | 35 |
| *Rhodopseudomonas palustris* | 7 | 10 | 10 | 27 |
| *Ruegeria* sp. TM1040 | 8 | 1 | 16 | 25 |
| *Salmonella enterica* Typhimurium | 13 | 7 | 8 | 28 |
| *Sulfitobacter* sp. EE-36 | 12 | 8 | 12 | 32 |
| *Xanthomonas campestris* ATCC 33913 | 6 | 6 | 3 | 15 |

Object 1. Table of promoter sequences for all species included in this work. PDF, 662 KB

### Conservation of Core Promoter Sequences

To survey trends in the promoter elements correlated with high activity in our dataset, we began by examining the core promoter elements. Sequences of the −10 and −35 core promoter hexamers are the most important determinants of promoter function (5) and we were interested in the relationship between conserved motifs and promoter output in each phylogenetic classes. We first categorized the promoters from each species into those with high, medium, and low outputs (promoter counts in each category listed in Table 4). *Ruegeria* sp. TM1040 is an exception as its library included mostly promoters with high or low expression and only one sequence expressed within a middle range (Object 1, Figure 2). We then used the core promoter elements from the promoters within each category to build sequence logos (Weblogos for all species available in Object 2, KpLogo probability logos for all species available in Object 3).

Object 2. Weblogos for all species included in this work. PDF, 728 KB

Object 3. KpLogos for all species included in this work. PDF, 850 KB

Because the housekeeping σ factors in Proteobacteria are highly conserved (15, 16), we expected the core hexamers of high activity promoters to also be conserved at most positions. Unsurprisingly, *E. coli* consensus motifs, 5′ – TATAAT – 3′ and 5′ – TTGACA – 3′ in the –10 and –35 region, respectively, were prominent in the logos of the high expression promoter groups (Figure 3). Within these motifs, certain positions are known to be more highly conserved than others due to specific interactions with σ^70^ during transcriptional initiation. The adenine at position –11 and the thymine at position –7 are thought to be highly conserved due to their stabilizing effect during DNA strand separation (24, 43). In our data, the thymine at position –7 was completely conserved in the high activity promoters across 14 species. A notable exception was *Xanthomonas campestris*, where the second highest expressing promoter contained a non-consensus cytosine in that position. The adenine at –11 is well conserved across the Beta- and Gammaproteobacteria but less so in the Alphaproteobacteria, where promoters screened in both *A. fabrum* and *R*. sp. TM1040 contain a guanine in that position. Indeed, a 5′ – AGTAAT – 3′ –10 element paired with a consensus –35 and extended – 10 element was the highest expressing promoter in *A. fabrum* (Object 1). The –12 position is important in fork junction recognition by σ^70^ (44, 45). The Alphaproteobacteria were also more likely to have a non-consensus nucleotide at positions –12, particularly in *R.* sp. TM1040, where three of the eight high expression promoters did not contain a thymine in this position. Sequence logos of the non-consensus –10 elements of the high expressing promoters in four Alphaproteobacteria reveal little conservation beyond the conserved –7 thymine (Figure 4). The 5′ – TTG – 3′ motif in the –35 element is also known to be highly conserved across species (24). As expected, our data shows that TTG is more conserved than the ACA motif at the 3′ end of the hexamer. Some Alpha- and Gammaproteobacteria did not have perfect consensus of this motif in the high expressing promoters (Figure 3). In ranking promoters from highest to lowest expressing, promoter 5 did not contain the TTG motif in *E. coli* and interestingly, the highest expressing promoter in *P. syringae* possessed a 5′ – ACG – 3′ motif in place of TTG, though both a consensus –10 and extended –10 element were present (Object 1). While these results were not statistically significant, possibly due to small datasets, it is nonetheless interesting to identify promoters lacking conserved nucleotides in positions that are thought to be highly conserved across species or necessary for transcriptional initiation. Though, we did not map transcriptional start sites experimentally, thus it is possible that alternate promoters are responsible for expression.

**Figure 3.**
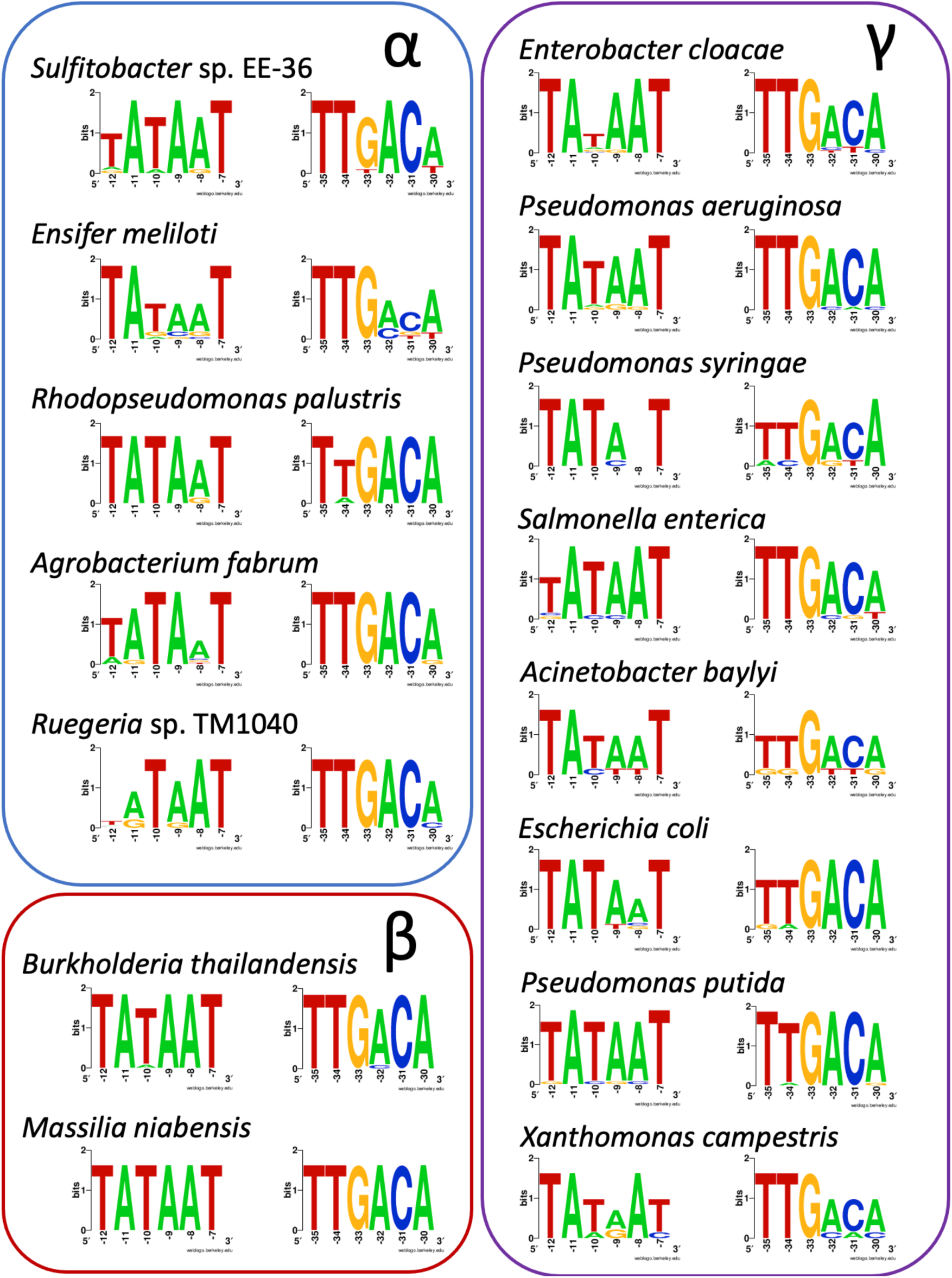
Weblogos of core hexamers from high expression promoter groups. Logos are grouped into species from the Gamma- (right), Alpha- (top left), and Betaproteobacteria (bottom left). The –10 hexamer is in the left column of each group and the –35 hexamer is at the right. Number of sequences used to make each logo listed in Table 4.

**Figure 4.**
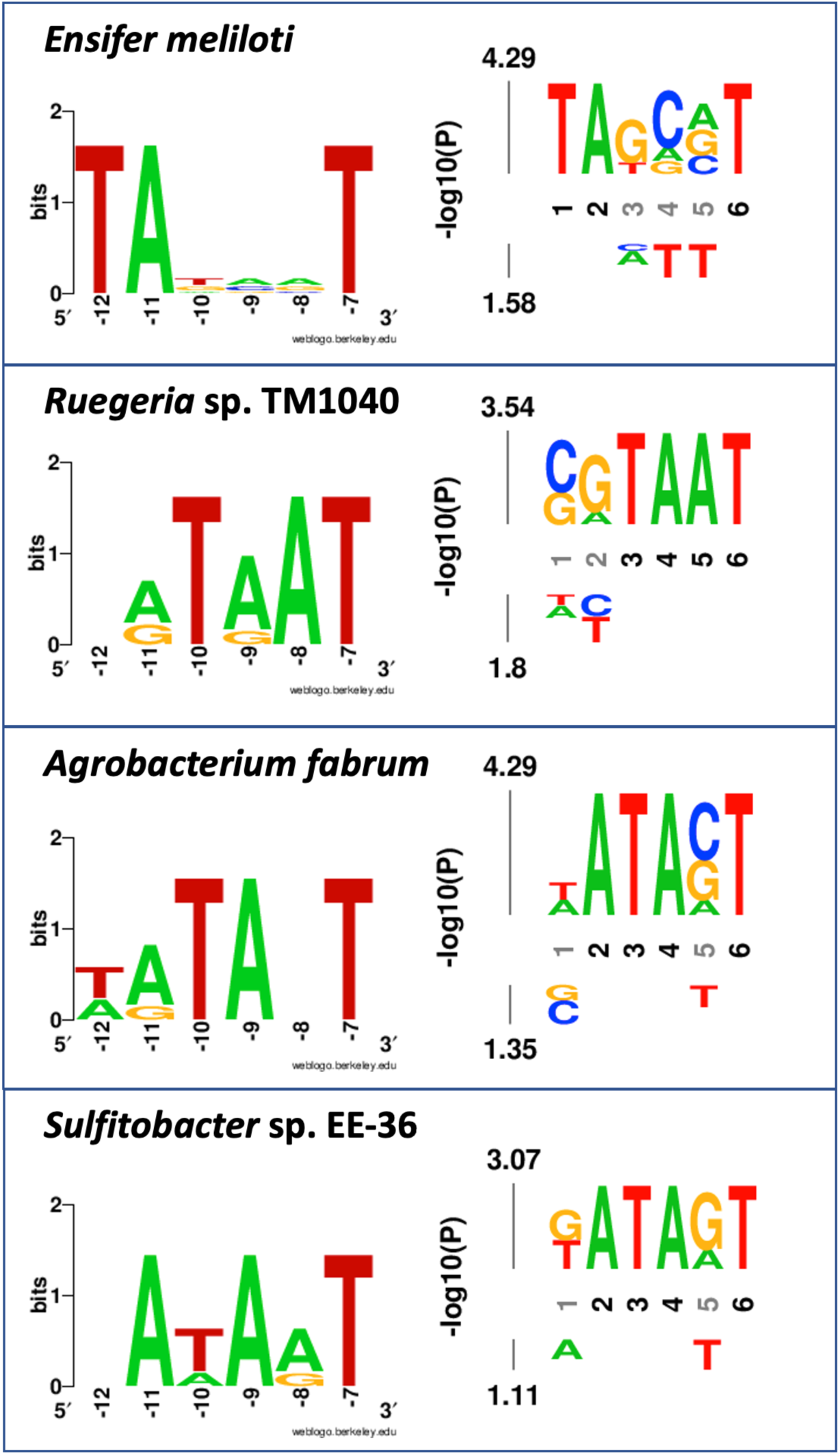
Logos of non-consensus –10 elements from high activity promoters in the Alphaproteobacteria. Left column are logos representing conservation in bits and right column are probability logos where residues are scaled relative to statistical significance of each residue at each position with −log10(p-value) on the y-axis. Apart from the conserved thymine at position –7, little conservation is apparent across this Proteobacterial family. *R. palustris* is excluded from these logos as only one high expressing promoter lacks a consensus –10 element.

The presence of consensus hexamers is not necessarily indicative of a highly active promoter. Even in our small datasets, two thirds of the species tested contained a promoter sequence with both consensus hexamers in the lowest expressing promoter category, including *E. coli*, *R. palustris*, *M. niabensis, B. thailandensis, E. meliloti, S. enterica, P. syringae, P. aeruginosa, S.* sp. EE-36, and *E. cloacae.* Over a quarter of the promoters in the low activity groups in *B. thailandensis, P. syringae*, and *R. palustris* contained –10 and –35 consensus sequences but were inactive regardless. This may be a result of RNAP holoenzyme binding these elements so tightly that it cannot effectively clear the promoter (10, 46, 47). Among promoters in the lowest expressing category that possessed one core consensus element, the –10 element was absent more frequently than the –35 element in most species. In our dataset, a consensus –10 element was absent in all inactive promoters in *A. baylyi*. Though *X. campestris* contained the smallest promoter library, our data shows that at least one consensus core element was required for moderate to high activity. The lack of a definitive association between the presence or absence of two consensus core elements and high or low activity, respectively, is consistent with previous studies (5, 48). This emphasizes the need for larger, more comprehensive datasets that can thoroughly explore interactions between core hexamers and other sequence elements that influence overall promoter output.

### The Extended –10 Element

The 5′ – TGn – 3′ motif at positions −15 to −13 is known as the extended –10 element and its presence has been shown to rescue activity from promoters with core hexamer sequences that deviate from consensus in *E. coli* and other Proteobacteria (47, 49). To investigate the effect of the extended –10 element in our promoter libraries, we analyzed sequences with and without the TGn motif in relation to the core promoter hexamers. Of the 15 species under study, *A. baylyi* and *E. meliloti* did not contain promoters with extended –10 elements in their characterized libraries. Across the remaining 13 species, 43 of the 416 promoters screened contained an extended –10 element. The motif was more often found in promoters with at least one non-consensus core promoter element than in sequences with the consensus. 20 of the 43 promoters were in the high expression categories across species. Of these, three promoters contained a consensus –35/extended –10 element pairing and two contained a consensus –10/extended –10 element pairing. In *A. fabrum* and *P. syringae*, these were the highest expressing promoters in their respective libraries, supporting evidence that an extended –10 element can compensate for weak core promoter elements (47). The remaining 23 promoters with an extended –10 element were paired with two consensus or two non-consensus core promoter elements.

While the extended –10 element is often discussed as a compensatory binding site for σ^70^ in the absence of two core promoter elements, this was not the case for many of the promoters in our dataset. 11 of the 20 high activity promoters with a TGn motif also contained consensus sequences for both core elements and in the remaining four promoters, neither core element was at consensus. For example, the eighth highest expressing promoter in *S. enterica* contained an extended –10 element and non-consensus 5′ – TACAAT – 3′ and 5′ – TTGCGA – 3′ motifs in the –10 and –35 core elements, respectively. Here, RFU measurements reached four orders of magnitude, the same as the highest expressing promoter of that library. In the ranking of highest to lowest activity promoters in *S. enterica*, the third promoter does not contain core consensus nor an extended –10 element (Object 1). Conversely, three highly active promoters containing an extended –10 element in *B. thailandensis* also possessed consensus core promoter hexamers. Though the –10 and –35 elements are at consensus, the distance between the elements may not be ideal for some species and as such, may benefit from additional binding sites like the extended –10 element (30, 37). Overall, the presence of an extended –10 element did not consistently yield a more highly active promoter. In fact, in *M. niabensis* and *R*. sp. TM1040, the TGn motif was only found in promoters with little to no detectable expression, even when the sequence contained one or both consensus hexamers (Object 1).

A recent study investigated the effect of multiple 5′ – TG – 3′ motifs centered at position –16. They found that while a single TG motif was present in the majority of strong promoters as TGn, tandem TG motifs dampened promoter activity (50). To investigate the effect of tandem TG motifs in our dataset, we took a subset of the spacer region from positions –19 to –14 and tallied the occurrence of 5′ – TGTG – 3′ and 5′ – TGTGTG – 3′ and we found one promoter that met this criteria. One of the lowest expressing promoters in *A. fabrum* contained 5′ – TGTGTG – 3′ at positions –19 to –-14. Because this promoter also contained non-consensus core hexamers that are a unique pairing in the *A. fabrum* promoter library, the contribution of the tandem TG motif to low expression is unclear and would be an interesting avenue for further study.

### The Spacer Region

While the spacer region between the two core promoter elements does not have a defined consensus sequence, its length and GC content are known to affect expression (51). Further, a recent study found a correlation between a specific region of the spacer element between positions –20 and –13 and expression in a screen of synthetic promoters in *E. coli* (5). Because promoters from the nonrepetitive promoter library have a set spacer length of 17 bp, we were interested in examining the GC content of the entire spacer as well as the –20 to –13 region to see how each affected activity and if the –20 to –13 region was a better indicator of promoter activity. We compared the GC content of the full spacer region from the high expression group to that from the low expression group for each species and visualized the data as violin plots (Figure 5). GC content in the high activity promoter group was not notably lower than that of the low expressing promoters, even in *E. coli*. Other studies have found a correlation between lower GC content and increased promoter activity and so this trend is somewhat surprising (5, 50). This could be due to the size of the dataset and the well-studied phenomenon of complex non-linear interactions among promoter elements in determining overall promoter function (5, 38, 47). The genome GC content of each species is another confounding factor (52). Though no obvious trends were apparent across the phylogenetic classes, some insight can be gained by examining more closely related species. In the Gammaproteobacteria, both *P. putida* and *P. syringae* have a cluster of low expressing promoters with a GC content of 47% while most of promoters in the high expression group have a GC content over 50%. In the Alphaproteobacteria, more than half of the low expression promoters had a GC content under 50% except for *R*. sp. TM1040. The only two species with a majority of high expressing promoters with spacer regions less than 50% GC content were *A. baylyi* and *B. thailandensis*. While *A. baylyi* has the lowest genome GC content of the 15 species at 40%, *B. thailandensis* is among those with the highest and so this difference is the most striking.

**Figure 5.**
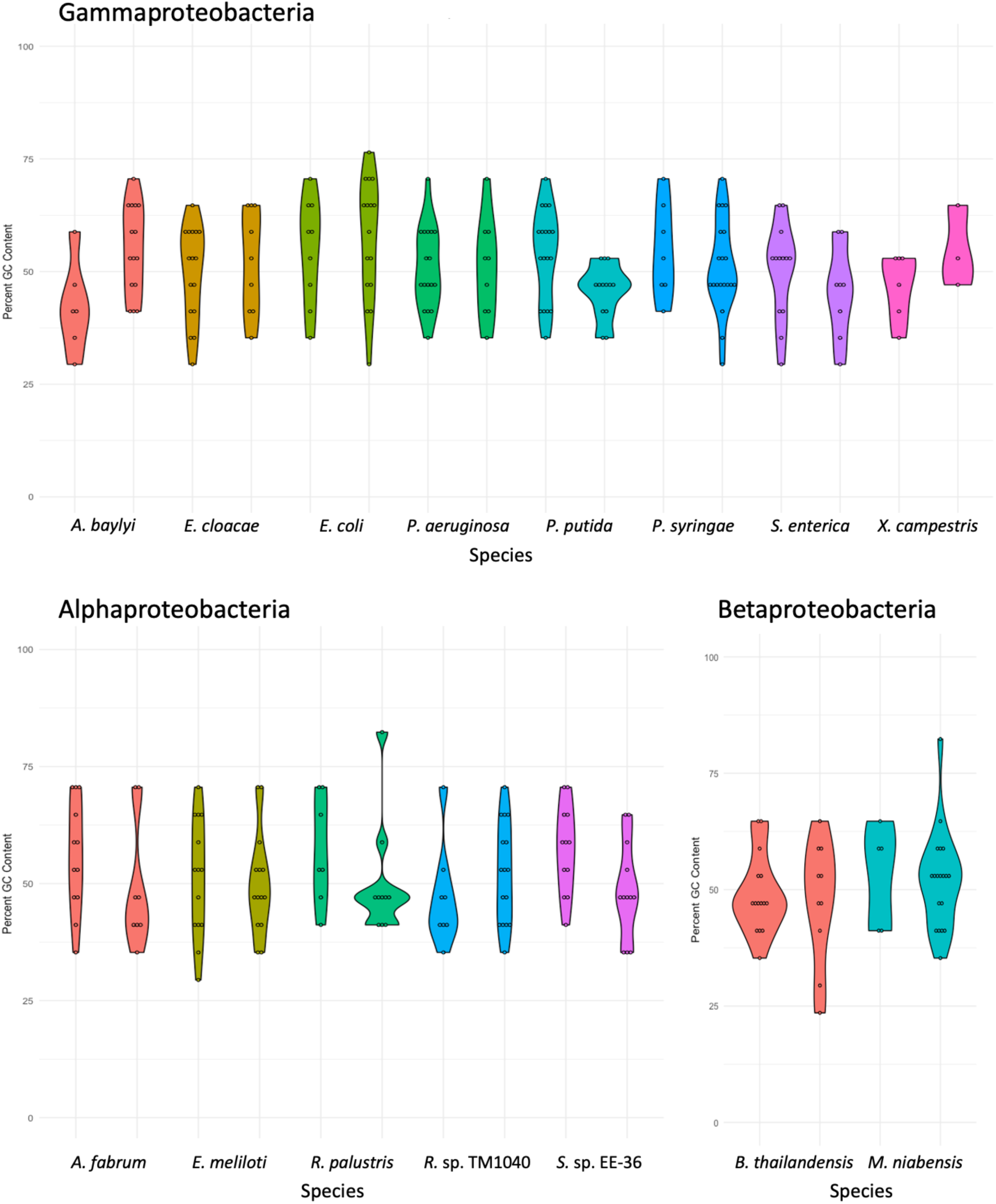
Violin plots of GC content of full spacer region for 15 species. Each colored pair of plots represents spread of GC content percentage in each species. In each pair of plots, the first represents data from the high expression promoter group and the second from the low expression group. Number and spread of datapoints used to make plots represented by open circles within each plot.

Next, we compared the GC content of the –20 to –13 spacer region across species. The –20 to –13 region in the spacer is proximal to the –10 element, which is where DNA strands are first separated during transcriptional initiation (21) and GC content at these positions may be more directly related to promoter activity. The –20 to –13 region in the spacer will be referred to as the proximal spacer for clarity. We analyzed these trends by generating another set of violin plots (Figure 6). For most promoter libraries across species, the rank order of promoters by GC content of the full spacer were similar to the rank order by GC content of the proximal spacer. While the lowest GC content of the full spacer region in a promoter across species was 18%, the proximal spacer GC percentage was as low as 12.5% for promoters in *A. baylyi*, *M. niabensis*, *S. enterica*, and *S*. sp. EE-36. For *S. enterica* and *A. baylyi*, at least one of these promoters was in the high activity category. Indeed, our results show that across most Gammaproteobacteria, the median GC content of the proximal spacer was lower than the median GC content of the full spacer in the high expressing promoters. Though, this was not true for *E. coli* or *P. aeruginosa*; in fact, *E. coli* had the highest median proximal spacer GC content of high activity promoters of all species screened at 75%. In comparison, the median GC content of the full spacer in this promoter group in *E. coli* was 59% and 54% for all promoters screened in *E. coli*.

**Figure 6.**
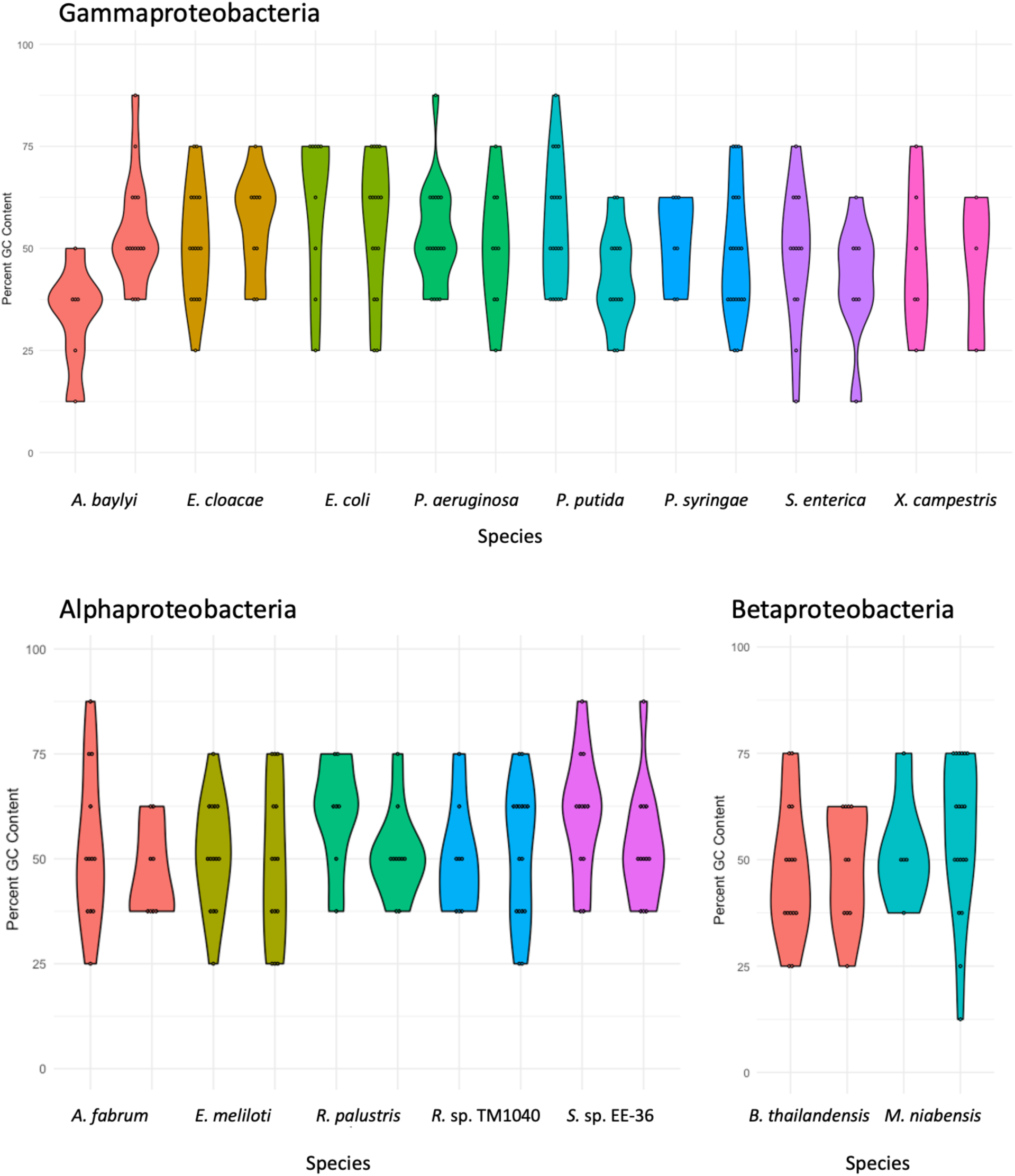
Violin plots of GC content of proximal spacer region for 15 species. Each colored pair of plots represents spread of GC content percentage in each species. In each pair of plots, the first represents data from the high expression promoter group and the second from the low expression group. Number and spread of datapoints used to make plots represented by open circles within each plot.

In some species, higher GC content in the full spacer and proximal spacer region were more likely to be correlated to promoters with very high expression. For both species of Betaproteobacteria, *R. palustris*, *R*. sp. TM1040, *E. coli,* and *X. campestris*, a promoter with 75% GC content in the proximal spacer was among the top three highest expressing promoters in that species (Object 1). In all but one case, this high GC spacer region was paired with two consensus hexamers. For some species, differences in GC content between high and low expression promoters is more exaggerated when the proximal spacer is analyzed rather than the full spacer, as in *A. baylyi* and *R. palustris*. Echoing the same trend, more proximal spacers in the high expression group have GC contents at or above 75% compared to GC contents of full spacers in most Gammaproteobacteria. As mentioned above, the datasets analyzed here are not large enough to outline definitive relationships between certain promoter characteristics and expression, though the high GC content of the spacer regions in high expressing promoters was somewhat surprising.

### Promoter Transferability

To investigate the transferability of our promoters between Proteobacteria, we chose a subset of the characterized promoters from our libraries to rescreen in another species. Specifically, a total of 47 promoters were selected from 4 species across the ɑ-, β-, and γ-proteobacteria, with 12 promoters from each species. These 12 promoters came from the 3 activity categories: high, mid, and low, with 4 promoters from each category. For each of the 4 strains, the set of 36 promoters not previously screened in that strain was screened for mRFP expression. For *S.* sp. EE-36, 11 promoters were rescreened, with 3 promoters in the low category. We refer to the species where promoter library expression data was collected as the source species and the species where the promoter was rescreened as the target species. The results of the promoter rescreening experiment are shown in Figures 7 and 8 and an accompanying table of promoter names and sequences is in Table 5.

**Figure 7.**
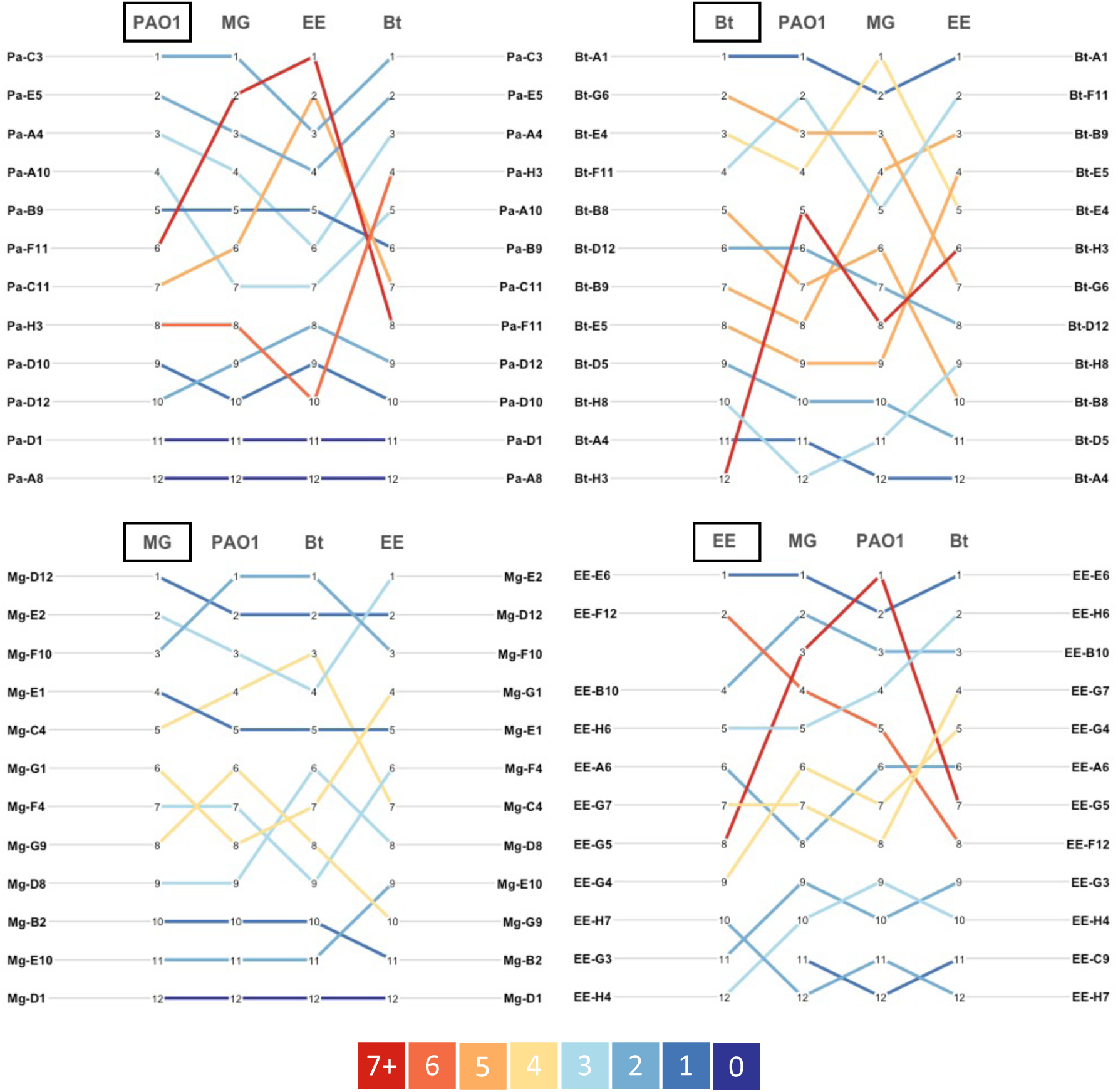
Slopegraphs of changing rank order of promoter activity from 47 promoters in four species. Graphs clockwise from top left display promoters selected from the *P. aeruginosa*, *B. thailandensis*, *S.* sp. EE-36, and *E. coli* datasets screened across the four species. Color legend indicates the maximum change in rank order of a promoter across the four species dataset within each slopegraph.

**Figure 8.**
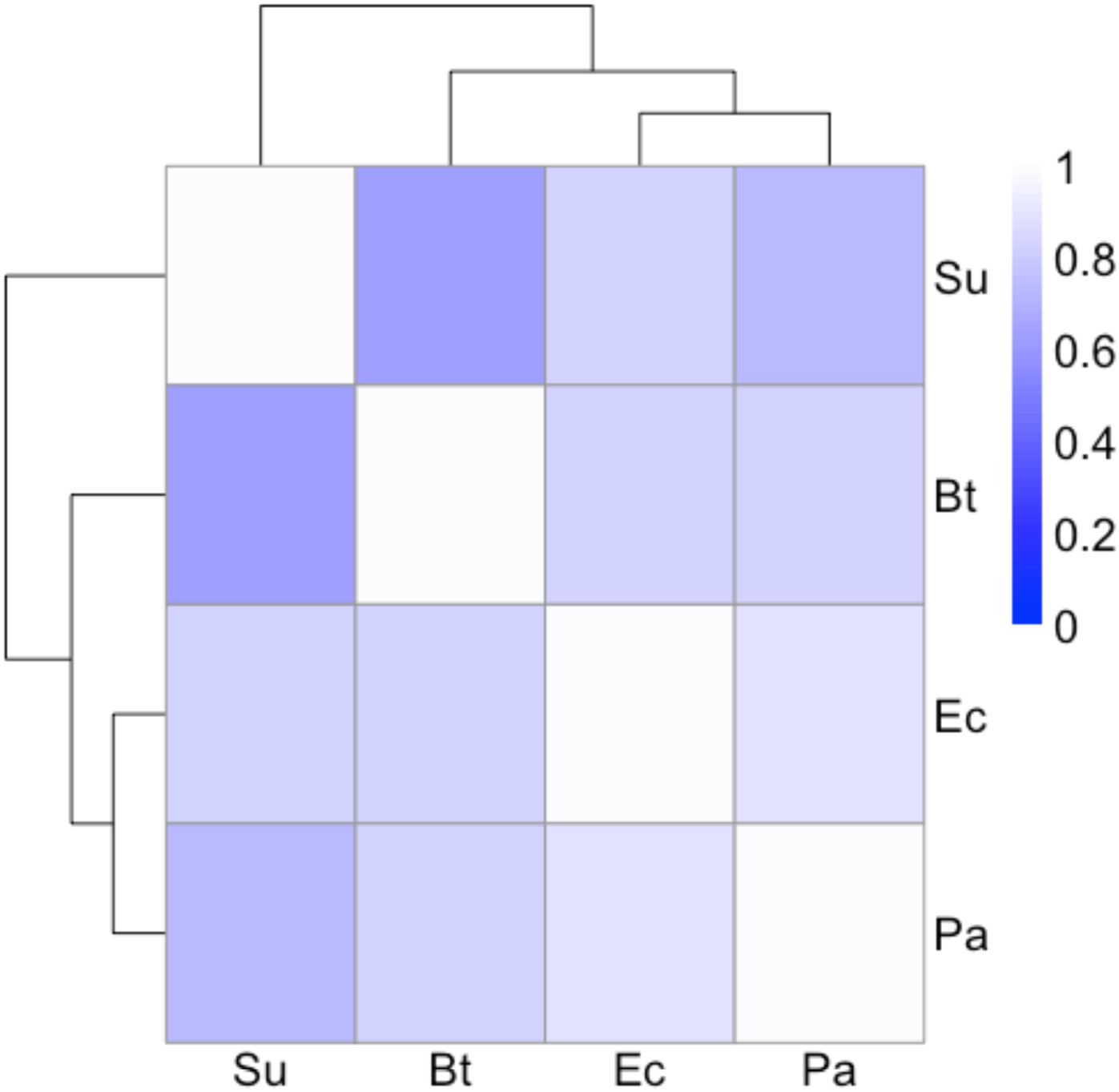
Heatmap of Spearman correlation analysis with clustering on rank order of 47 promoters in each of four species: Su: *S.* sp. EE-36, Bt: *B. thailandensis*, Ec: *E. coli*, Pa: *P. aeruginosa*.

**Figure 9.**
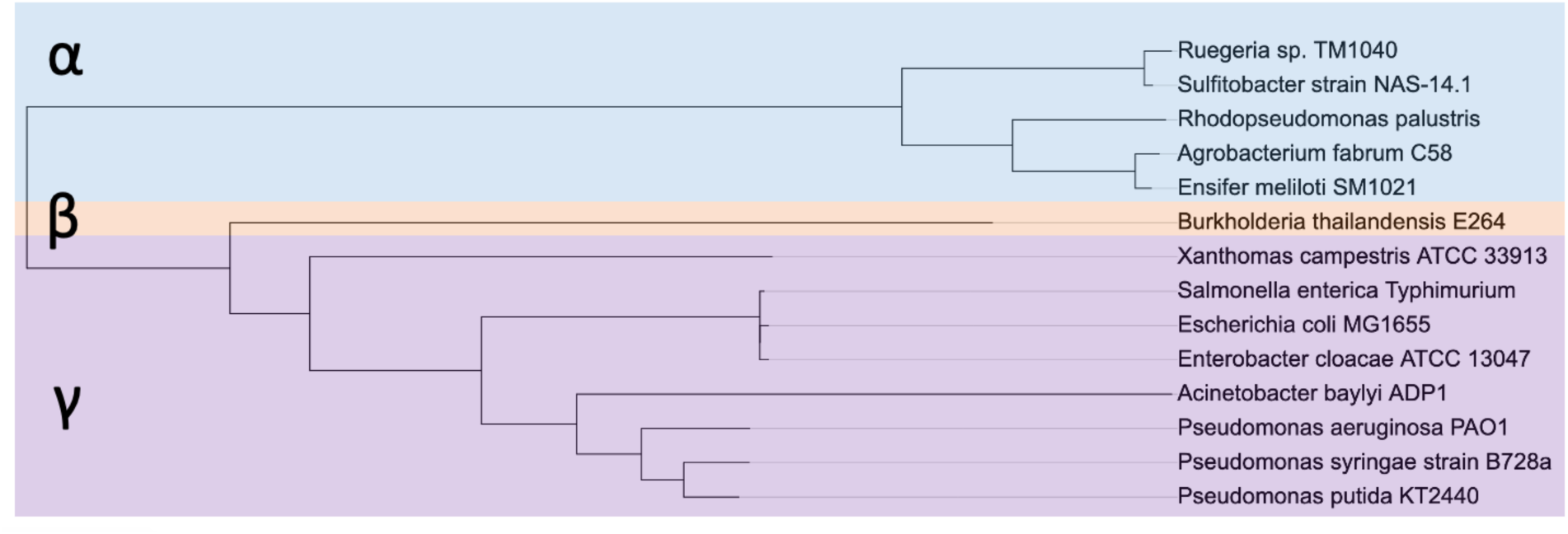
Phylogenetic tree constructed from housekeeping σ factor sequences. Tree represents the phylogenetic relationship of the housekeeping σ factors for Alpha-, Beta-, and Gammaproteobacteria species included in this work except for M. niabensis as no RpoD protein sequence is available for this species. For S. sp. EE-36, the closely related S. strain NAS-14.1 is used in place.

**Table 5:**
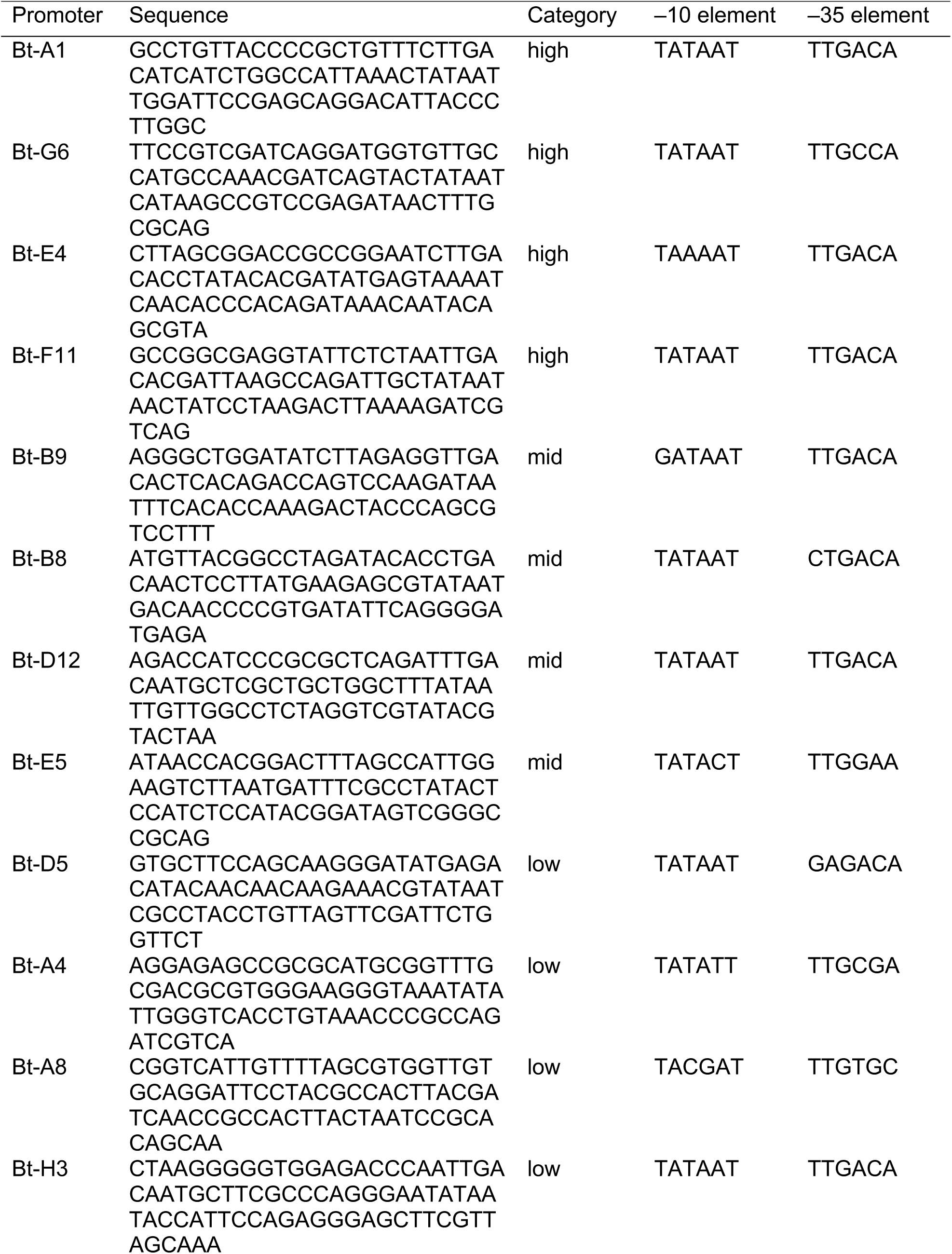
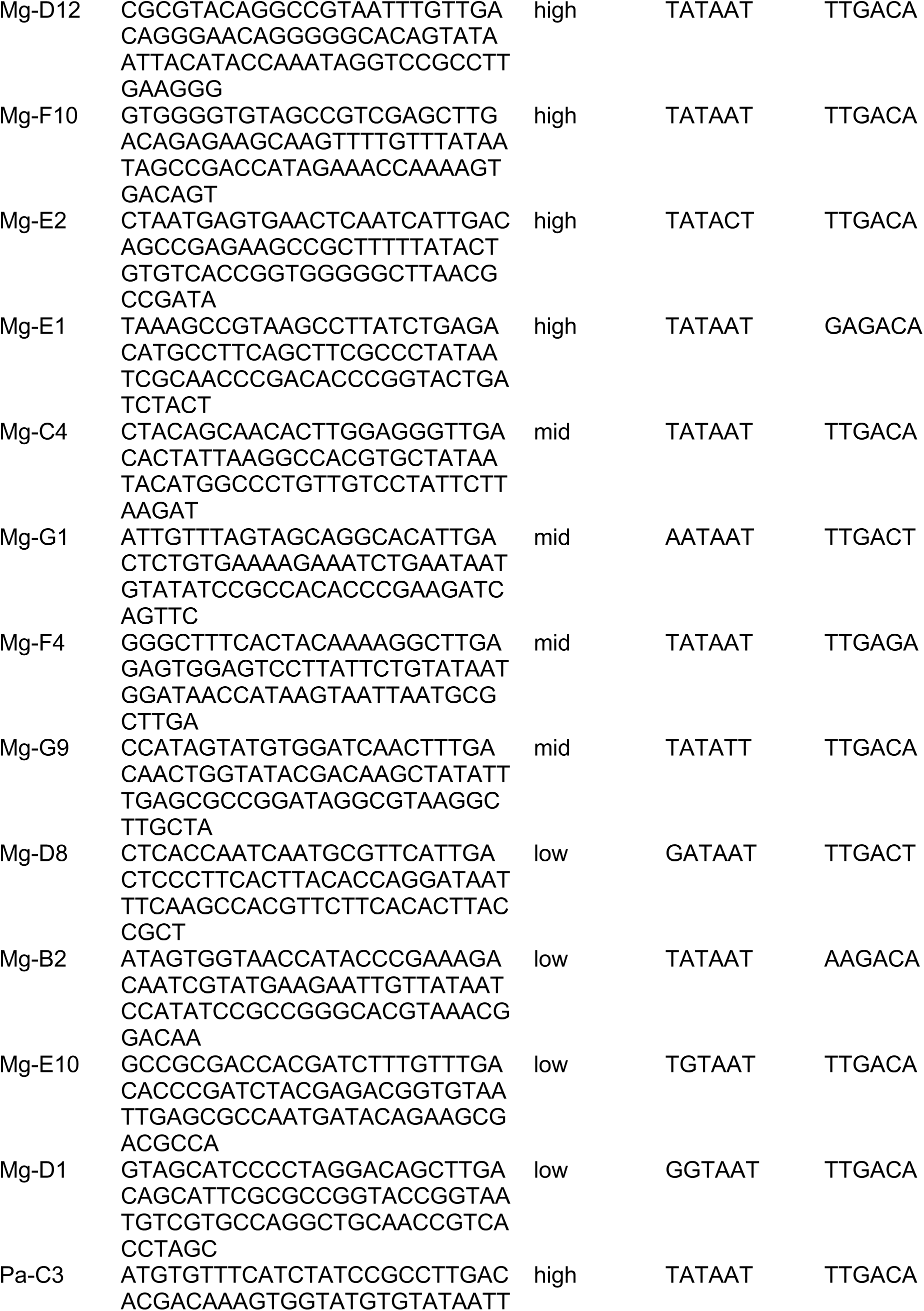
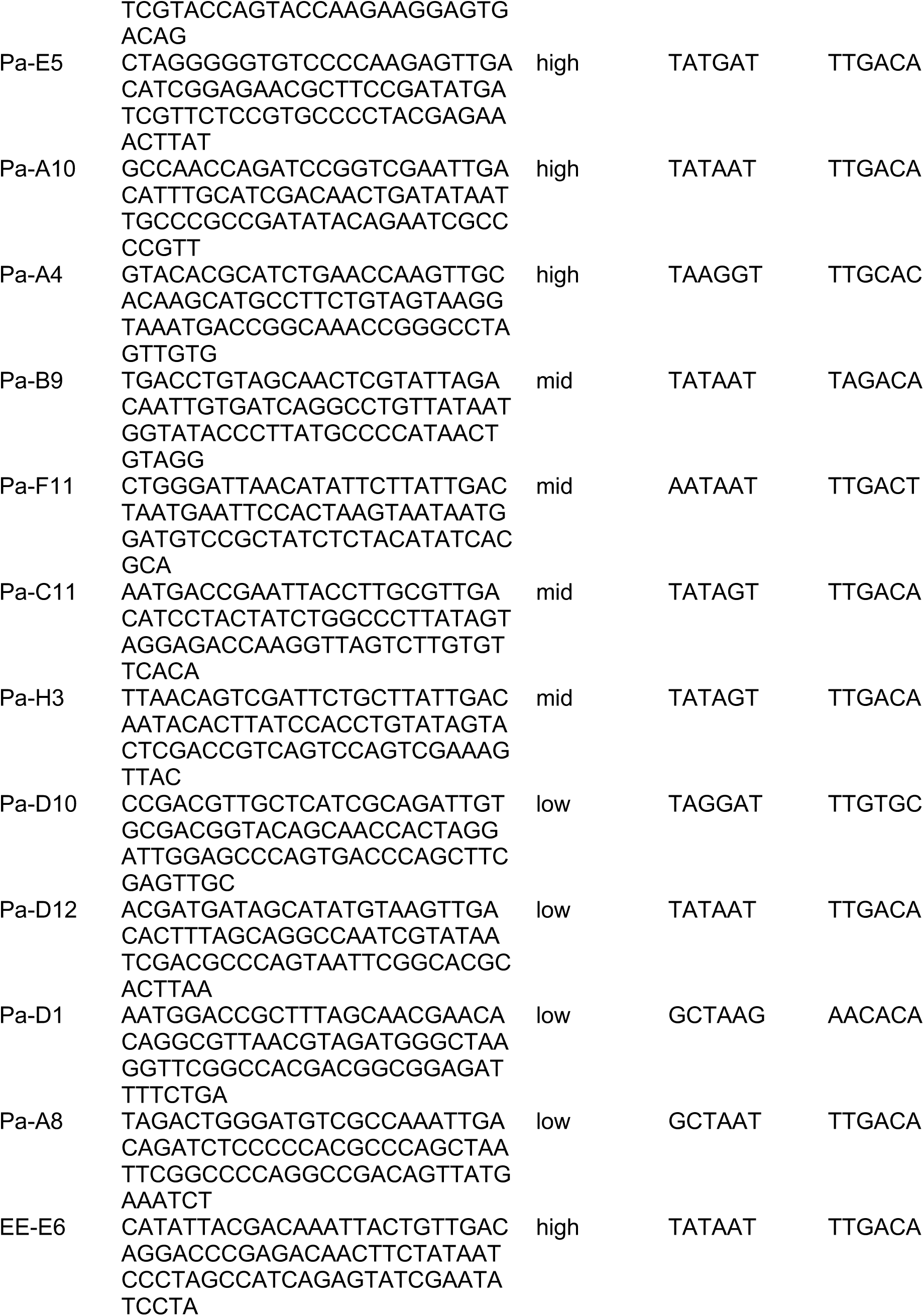
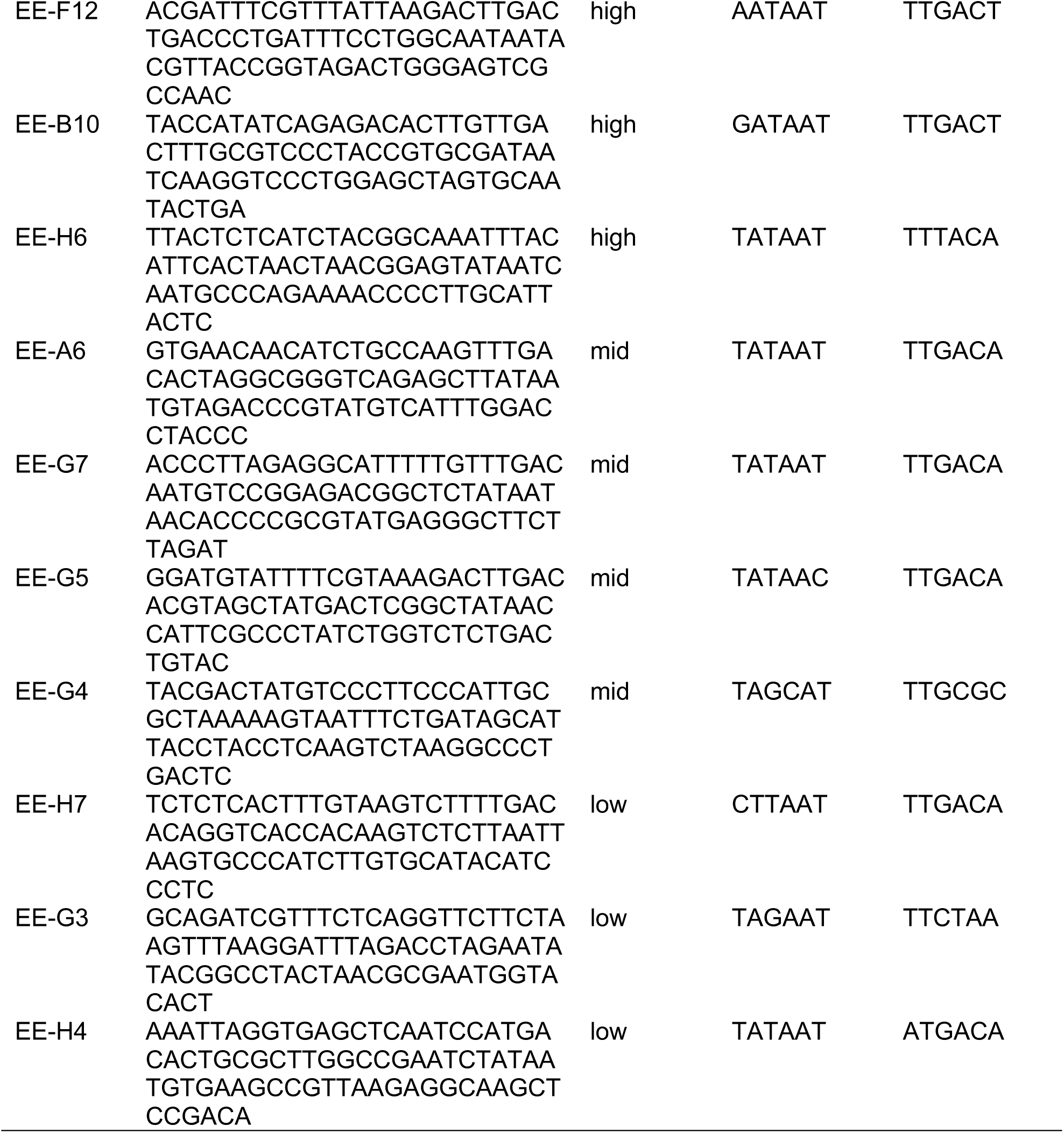
Promoter Transferability Sequences.

In three out of the four source species’ promoter sets, there was a notable shift in activity rank order, with a maximal change of at least seven ranks observed across the species. In the datasets of source species *P. aeruginosa* and *S.* sp. EE-36, the promoter that exhibited this significant shift held the top rank one species. Specifically, the promoter Pa-F11 was rank 1 in *S.* sp. EE-36 and experienced a drop of seven ranks when measured in *B. thailandensis*, while promoter EE-G5 was rank 1 in *P. aeruginosa* and saw a similar drop when measured in *S.* sp. EE-36. In the *B. thailandensis* dataset, a promoter ranked 12th in B. thailandensis ascended 7 positions when measured in *P. aeruginosa.* These large rank transitions occurred when promoters were measured in species in different phylogenetic classes.

To test our hypothesis that a greater phylogenetic distance between species corresponds to more significant alterations in the rank order of promoter activity among those species, we compared the summed rank order changes between source and target species pairs within each dataset (Table 6). We found that the largest cumulative rank order change occurred when Pa promoters were measured in *P. aeruginosa* compared to measurements in *S.* sp. EE-36, with a total of 24 rank order changes. Moreover, the dataset from the source species *S.* sp. EE-36 exhibits the highest cumulative rank order change when these changes are aggregated across all pairings of source and target species within the dataset. Alphaproteobacteria are more distantly related to Gammaproteobacteria than Betaproteobacteria (53), so this is consistent with our hypothesis. This is further validated by a Spearman correlation analysis conducted on the ranked data, where clustering was consistent with the clustering derived from a phylogenetic analysis (Figure 8). A cumulative rank order change of 10 was the lowest measured across source species datasets. This occurred when Mg promoters were measured in *E. coli* compared to measurements in *P. aeruginosa*, and, intriguingly, was also observed when Pa promoters were measured in *P. aeruginosa* compared to *B. thailandensis* and when Bt promoters were measured in *B. thailandensis* compared to *P. aeruginosa*. The comparatively similar promoter expression profiles observed between *P. aeruginosa* and *B. thailandensis* could be attributed to the fact that both species exhibit a GC content of 68%.

**Table 6:** Cumulative rank order changes in pairwise species.

| Source species dataset | Species pairing | Total sum of rank order changes |
| --- | --- | --- |
| PAO1 | *P. aeruginosa* and *E. coli* | 12 |
| PAO1 | *P. aeruginosa* and *S. sp. EE-36* | 24 |
| PAO1 | *P. aeruginosa* and *B. thailandensis* | 10 |
| MG | *E. coli* and *P. aeruginosa* | 10 |
| MG | *E. coli* and *S. sp. EE-36* | 14 |
| MG | *E. coli* and *B. thailandensis* | 14 |
| Bt | *B. thailandensis* and *P. aeruginosa* | 10 |
| Bt | *B. thailandensis* and *S. sp. EE-36* | 22 |
| Bt | *B. thailandensis* and *E. coli* | 14 |
| EE | *S. sp. EE-36* and *P. aeruginosa* | 20 |
| EE | *S. sp. EE-36* and *B. thailandensis* | 22 |
| EE | *S. sp. EE-36* and *E. coli* | 14 |

In agreement with our promoter library screening data, the presence of *E. coli* consensus sequences in the –10 and –35 hexamers did not consistently correlate with high promoter activity. The most striking example of this is with promoter Bt-H3, which was the lowest ranked promoter in *B. thailandensis* and ranked within the mid category in *P. aeruginosa*, *E. coli*, and *S.* sp. EE-36, but contained perfect core promoter motifs. Promoter Pa-D12 also contained perfect core sequences but remained in the low category in two of the three target species. Conversely, Pa-F11 is the rank 1 promoter in *S.* sp. EE-36 while containing a single mutation in both the –10 and –35 hexamers. Core hexamers are not the only elements that determine promoter activity level, and we have considered the GC content of the spacer region in addition to the presence of an extended –10 element, but no clear trends emerge. For example, high GC content in the spacer region of the promoter is associated with lower activity (50), but we did not observe this trend consistently across species.

Surprisingly, though the promoter with the highest activity from each source species remained in the high category across the three target species, the rank 2, 3, and 4 promoters in each source species did not. While these categories are somewhat arbitrary, combined with inconsistencies in the relationship between core promoter hexamers and promoter activity, this emphasizes the unexpected problems that can emerge when a promoter is presumed to have inherent high activity based on its performance in a single species. Consistency in rank order activity across species was most apparent across promoters in the low category, with the exception of Bt-H3. This suggests that these promoter sequences may have features or a combination of features in common that is less conducive to RNAP promoter binding and escape.

## Discussion

In this study, we established a toolbox of synthetic constitutive promoters characterized in 15 diverse species of Proteobacteria. The toolbox includes libraries of 15-43 promoters tested in each species and promoter activity spans 3-5 orders of magnitude in each library. From this data, we surveyed trends in core promoter sequences, the GC content of the spacer region, and the presence or absence of an extended –10 element in high expressing promoters. Though our dataset was too small to draw significant conclusions, we noted characteristics of promoters that behaved in unexpected ways and compared high activity promoters across species in the same Proteobacterial class. We then tested the transferability of some promoters by selecting a subset for rescreening in another species. In this study, it was observed that the majority of rescreened promoters underwent a shift in their rank order of activity by more than two ranks across each source species dataset. Notably, a promoter transitioned seven ranks from one species to another in three out of the four datasets. Data gathered from these experiments highlight the variable activity of promoters across different host, even when hosts are closely related species. Promoter transferability is valuable in its own right and complements the promoter toolbox with cross-species promoter characterizing. This work represents the most comprehensive characterization of a promoter toolbox across Proteobacterial species to date.

Prior to this work, there were few established collections of constitutive promoters for use in non-model bacteria (54, 55). Of those, many consist of endogenous promoters, identified and reused in the same host. Using native promoters in synthetic circuits can be problematic as they are more likely to contain cryptic regulatory elements and alternative transcriptional start sites, making them host context-dependent and not orthogonal genetic parts (54, 56). For these reasons, we chose to test synthetic promoter libraries as they offer more reliable and reproducible expression (57). Constitutive promoter libraries have proven to be valuable for the construction and testing of genetic systems as they provide options for fine-tuning expression levels. For example, the Anderson promoter library is among the most popular collections of synthetic constitutive promoters and includes 19 promoters spanning a gradient of expression levels in *E. coli* (58). Though the collection was developed for use in *E. coli*, it has been utilized in diverse species to optimize expression of transcriptional regulators, control the expression of an sgRNA, and expand toolkits for species where few characterized promoters are available (59–63). This exemplifies the demand for characterized genetic parts in non-model species as these promoters are often chosen due to a lack of alternatives (63). The promoter libraries tested in this work are made available here to meet this need, providing sequence and output of at least 15 promoters in 15 species spanning the Alpha-, Beta-, and Gammaproteobacteria.

More recently, synthetic promoter libraries have been used to study the relationship between promoter sequence and function. In a landmark study by Urtecho et al., the authors employed a massively parallel reporter assay to explore the transcriptional activity of over 10,000 promoter variants in *E. coli* (5). Their library consisted of every combination of a set of discrete promoter elements, including eight – 35 elements, eight –10 elements, three UP elements, eight spacer regions, and eight background sequences. From their expression data, they were able to train a statistical model to predict promoter strength and describe more complex interactions between elements that increased or hampered activity. Another study screened over 14,000 promoters to predict site-specific transcriptional rates for σ^70^-dependent promoters in *E. coli*, with over 100 variants each of the UP element, –10 and –35 core elements, spacer, extended –10 element, discriminator, and initially transcribed region (48). While our goal was different and focused on characterizing promoter activity in non-model species, we surveyed the results of our screens to find notable trends and promoters with unexpected results. Unlike the screens mentioned above, our datasets were too small to draw statistically significant conclusions or definitively identify markers of high promoter function in the Proteobacteria; instead, our work epitomizes the complex and non-linear relationships among elements that contribute to promoter function. For example, in 11 of the 15 species included in our screen, at least one promoter in the high activity category did not possess a consensus core promoter element and in 10 of the 15 species, we found at least one inactive promoter with both consensus hexamers. These consensus sequences are based on research in *E. coli* but they are likely similar in other Proteobacteria based on existing research (30, 31, 33) and the high conservation among housekeeping σ factors (25, 26). Though, core sequences are only part of a larger picture when considering the determinants of a highly active promoter.

While not done to the scale of related work in *E. coli* (5, 38), our work still identified some interesting and contradicting relationships between sequence and expression that could be further explored in more systematic screens of promoter activity. One unexpected trend in our dataset was the high GC content of the spacer regions of highly active promoters. In larger studies of promoter function in *E. coli*, GC content is negatively correlated with the level of promoter expression (5) but most work investigating promoters in other Proteobacteria has focused on the length of the spacer region and not its sequence composition (31, 33). A recent study investigating spacer elements has suggested that the GC content of the spacer region plays a crucial role in DNA supercoiling sensitivity, impacting the timing of expression and potentially the phasing of the –10 and –35 core promoter regions on the face of DNA (50). The effect of spacer region length and sequence on overall promoter output provides another avenue to further explore promoter function in non-model species. The effect of an extended –10 element could be better explored in a larger dataset as well. Similar to its function in *E. coli*, there is evidence that this motif functions to stabilize the RNAP holoenzyme and compensate for weak core element binding in other Proteobacteria as well (47). The contribution of the extended –10 element to overall promoter function has not been thoroughly explored outside of *E. coli* (5, 38) and would further our understanding of promoter function across species.

In this study, we rescreened a subset of promoters in three additional species to evaluate variations in activity when the same genetic part is moved into a new host. Cross-species promoter characterization is valuable as research has shown promoter behavior is not consistent even between closely related species (39, 64). This may be due in part to transcriptional laxity, which refers to the capacity to efficiently transcribe from mutated endogenous promoters or horizontally transferred DNA (37). Some species, particularly the Alphaproteobacteria, have higher transcriptional laxity and as such, are able to utilize a larger range of sequences effectively as promoters (31, 37). In our experiments, the highest activity promoters screened in *P. aeruginosa* had expression levels that were an order of magnitude higher than high activity promoters in all other species (Object 1). *P. aeruginosa* is an opportunistic pathogen with a complex metabolism that enables it to adapt to free-living and pathogenic lifestyles (65); hence, an increased transcriptional laxity in this species would be evolutionarily beneficial. The same may be true of *A. fabrum*, *P. putida*, and other metabolically diverse species, but our dataset is not large enough to capture these trends. Given a larger dataset, we hypothesize that species that are opportunistic pathogens or possess diverse metabolisms would have a high degree of transcriptional laxity and an increased likelihood of effectively using a given promoter. It would be interesting to systematically re-screen promoters through this lens, comparing levels of transcriptional laxity to the lifestyles and metabolisms of diverse Proteobacteria.

Overall, we characterized libraries of constitutive promoters in 15 diverse species of Proteobacteria and further characterized a subset of these promoters across additional hosts to test the transferability of the genetic parts. Though our dataset was too small to draw conclusions, we note interesting and unexpected sequence-function relationships that can be further explored in a more comprehensive study. Our work highlights complexities of determining promoter activity from sequence across species despite the high conservation of the housekeeping σ factors in Proteobacteria. Indeed, a phylogenetic tree made from the protein sequences of housekeeping σ factors in the 15 species studied here is consistent with that that of a tree constructed from a 16S rRNA alignment (Figure 1, Figure 10). In a multiple sequence alignment of housekeeping σ factors in these species, σD2 is 100% conserved and σD4 is less but still highly conserved(21, 25, 26). The discrepancy between the high conservation of σD2 and σD4 and variability in the sequences of highly active promoters suggests greater complexity in the sequence determinants of a strong promoter than proximity of core elements to the canonical *E. coli* consensus. In characterizing promoters across species and identifying transferrable and non-transferrable sequences, we have contributed to the body of work focused on promoter sequence-function relationships. The study of promoters in different species and how they behave when transferred between hosts is highly valuable to the field and will accelerate the study and engineering of previously unexplored bacteria.

## Notes

https://original-ufdc.uflib.ufl.edu/IR00011835/00001

https://original-ufdc.uflib.ufl.edu/IR00011867/00001

https://original-ufdc.uflib.ufl.edu/IR00011837/00001

### Competing Interest Statement

The authors have declared no competing interest.

